# The Brain as Actor and Evaluator: Distinct Neural Codes for Timing and Self-Evaluation of Timing Errors Revealed by Transformer-Based Decoding

**DOI:** 10.64898/2026.03.24.714009

**Authors:** Sena N. Bilgin, Dunia Giomo, Urfan Mustafali, Tadeusz W. Kononowicz

## Abstract

Metacognitive self-evaluation, the capacity to assess one’s own performance without external feedback, is fundamental to adaptive cognition. A central unresolved question in metacognition is whether self-evaluation reflects a direct readout of the neural signals supporting first-order performance, or whether it depends on partially distinct higher-order representations. Unusual approach of monitoring actions in a continuous dimension, such as time, is particularly useful because errors are graded rather than binary, allowing metacognition to be treated as a quantitative estimate of error magnitude. Temporal error monitoring (TEM), in which individuals judge the accuracy of self-generated time intervals using only internal signals, provides a tractable framework for isolating metacognitive evaluation from external correction. We asked whether TEM engages the same neural mechanisms as first-order timing performance, the execution and control of the timed action itself, or recruits at least partially distinct processes involving additional evaluative computation. We decode single-trial EEG using a Vision Transformer applied to PCA-optimized θ, α, and β signals recorded during both monitoring of motor timing. First-order timing was decodable from each frequency band independently, whereas TEM required simultaneous integration across all three bands. Individuals for whom TEM neural states were decoded with greater accuracy demonstrated stronger behavioral error-monitoring precision. These findings establish TEM as a neurally distinct re-representation of timing performance, demonstrating that metacognitive self-evaluation requires evaluative computations beyond those supporting motor action execution.

## 1) Introduction

The capacity to monitor and evaluate one’s own cognitive processes, metacognition, is fundamental to adaptive behaviours across species.^1,2^ Temporal error monitoring (TEM) is one well-defined instance of this capacity: the ability to assess the accuracy of self-generated time intervals without external feedback. By isolating self-evaluation from exogenous cues, TEM offers a powerful framework for probing the neural basis of self-monitoring. A central question, however, remains unresolved: do metacognitive evaluations arise from the same decision variable as first-order performance, or do they at least partially engage distinct processes supported by additional neural representations?^3,4^

Timing tasks are particularly well-suited to address this question: unlike most cognitive domains, they require reliance exclusively on internal representations, with no external cues available. In motor timing paradigms, an agent generates a time interval by initiating it with one action and terminating it with a second when the target duration is judged to have elapsed. The discrepancy between intended and produced duration constitutes the error signal; estimating its magnitude and direction in the absence of feedback is the core requirement of TEM. Reliable TEM performance across both humans^5-10^ and rats^11^confirms that internally generated signals alone are sufficient to sustain metacognitive evaluation.

Accounts differ, however, in what they propose about the architecture that supports it. Single-process accounts hold that confidence and performance share a common decision variable, such that metacognitive judgements directly reflect task-execution signals with no additional processing required.^12^ Hierarchical models, such as the Bayesian framework of Fleming and Daw,^13^ posit a functionally distinct metacognitive stage; yet this stage still derives its content directly from first-order signals, computing confidence from their strength or uncertainty, making metacognition temporally later but not neurally distinct. Higher-Order Representation (HOR) theory draws a sharper distinction: conscious awareness of a mental state requires a higher-order representation that takes the first-order state as its object, a genuine re-representation rather than a downstream derivation from first-order signals.^14^ Neural evidence supports this view, implicating specialized prefrontal circuits in higher-order representations that cannot be reduced to first-order activity.^3,15^

Each of these positions generates a distinct prediction for TEM. Drift-diffusion models hold that error evaluation arises from the same accumulator governing interval production, implying shared neural substrates and a tight trial-by-trial coupling between timing performance and TEM precision.^12,16^ Temporal Metacognition, by contrast, views error evaluation as an active inferential process in which the brain decodes and re-represents internal duration signals at a higher level, not a mere reflection of the timing signal but a higher-order operation that draws on, yet goes beyond, first-order timing information, consistent with HOR theory.^4,14^

Distinguishing between these predictions, however, requires evidence that condition-averaged analyses cannot provide. Prior work linked beta-band oscillations to timing accuracy^17,18^ and metacognitive sensitivity,^4^ alpha oscillations to clock-speed modulation^19^ and temporal expectation,^20^ and frontal midline theta to prediction errors scaling with error magnitude.^21^ Whether these mechanisms are shared between first-order timing and metacognitive evaluation, or whether metacognition requires their coordinated interaction, cannot be resolved from averaged data alone.

To address this, we applied a Vision Transformer decoding architecture^22,23^ to single-trial EEG recorded during a motor timing task with no external feedback. First-order timing decodability did not predict trial-by-trial TEM precision, challenging drift-diffusion accounts,^12,16^ while metacognitive state decodability predicted TEM precision independently of timing performance. These findings argue against single-process and hierarchical accounts and support HOR theory, establishing metacognitive evaluation as an at least partially distinct re-representation of the brain’s self-evaluative architecture.

## 2) Methods

### 2.1) Subjects

A total of 29 participants (17 male, 12 female; age range: 18–36 years; M = 24.10, SD = 4.28) were recruited. All participants reported normal or corrected-to-normal vision. Written informed consent was obtained from each participant before data collection. All procedures were approved by the Research Ethics Committee at the Institute of Psychology of the Polish Academy of Sciences (reference: IP.403.3.2022) and conducted in accordance with the Declaration of Helsinki.^24^

### 2.2) Apparatus

#### Stimulus presentation

Subjects sat at approximately 55–60 cm distance from the computer monitor. Visual stimuli, task instructions, and feedback were presented using the Psychophysics Toolbox^25^ implemented in MATLAB. Behavioral responses were recorded via keyboard and mouse.

#### EEG recording

Electroencephalographic data were acquired using a 64-channel actiCAP snap cap (Easycap) with gel-based electrodes and an ActiCHamp amplifier (Brain Products). The EEG signal was sampled at 500 Hz. Electrode impedances were maintained below 10 kΩ throughout each recording session.

#### Data processing and analysis

All preprocessing and analysis pipelines were implemented in Python (v3.12.5). EEG signal processing was performed using MNE-Python (v1.9.0), and deep learning models were built in PyTorch (v2.5.1) within the TorchEEG framework. Code is available as described in the *Code Availabilit*y statement.

### 2.3) EEG Experiment Procedure

This study analyzed a subset of data (approximately 25%) from a larger study combining EEG and transcranial direct current stimulation (tDCS). EEG was recorded from a custom 64-channel cap with openings at Fpz and Fz, where neurostimulation electrodes were placed (anode at Fpz, cathode at Fz). Data were therefore recorded from 63 channels, with those two sites excluded.

The full experiment included a tDCS session and a Sham session, conducted on separate days to reduce cognitive fatigue and potential tDCS aftereffects (see Supp. Mat. Table 1). Each session comprised two blocks of approximately 20 minutes each. The first block was either a tDCS or Sham block, with order counterbalanced across participants. During the tDCS block, stimulation was delivered continuously at 1 mA, whereas in the Sham block, current was applied only briefly at onset before being discontinued to mimic the sensation of stimulation.

The second block consisted of an EEG-only recording. Because tDCS can induce aftereffects that persist beyond the stimulation period, only the EEG-only block of the Sham session yields uncontaminated measures of brain activity. Accordingly, all analyses in this study were restricted to this block. A 5-minute break was provided between blocks to minimize fatigue. Event markers were transmitted from MATLAB to the EEG system to label critical time points within each trial, enabling the extraction of epochs corresponding to pre-event, event, and post-event periods relevant to TEM.

Within this EEG-only recording, participants performed both time production (TP) and TEM tasks (Figure 1). Specifically, they were required to reproduce a 2-second target duration using a computer mouse. The target duration was introduced and practiced during a structured training phase to ensure that participants developed a stable internal representation of the interval.

**Figure 1.**
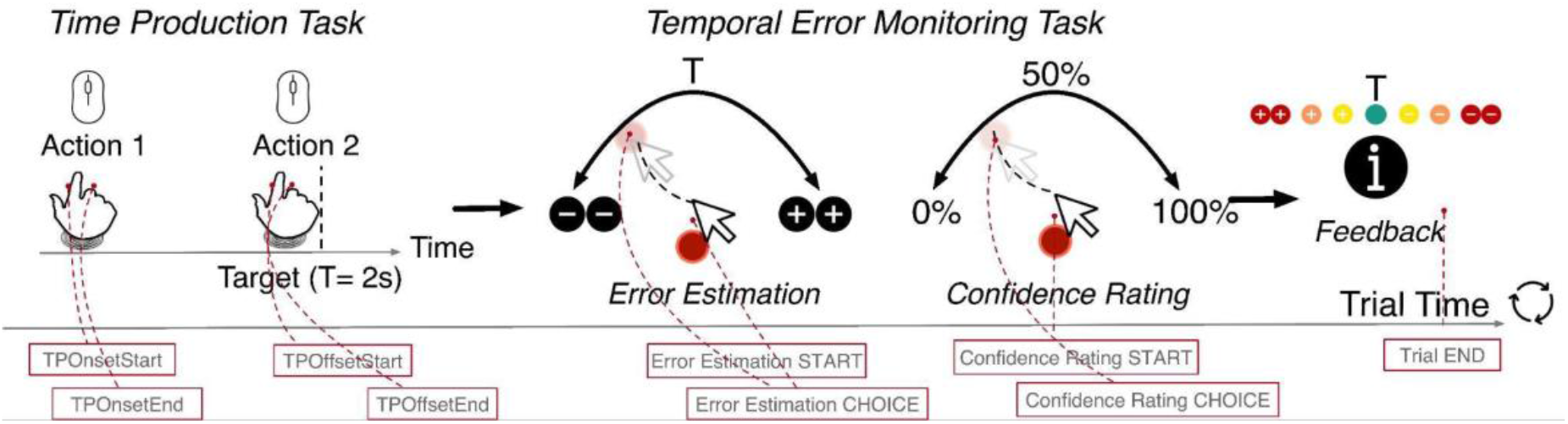
Experimental procedure. Participants reproduced a 2-second target interval by pressing a mouse button to mark its onset and offset. Following each reproduction, they provided two metacognitive judgments: a temporal error estimate indicating the perceived direction and magnitude of their timing error, and a confidence rating reflecting certainty in that estimate. Behavioral measures extracted from each trial included produced duration, temporal error estimation, and TEM confidence. The Metacognitive Inference (MI) index quantified trial-by-trial correspondence between objective timing performance and subjective error reports. Together, these measures enabled independent assessment of timing precision, metacognitive sensitivity, and confidence calibration.

#### a) Training Phase

Training was structured into three successive phases; each designed to progressively establish and refine the components of the task.

In the first phase, participants were familiarized with the 2-second target duration through 10 visually guided trials. Each trial began with a white fixation cross, which turned green to signal the onset of the interval; after 2 seconds, the green cross disappeared, followed by a randomly selected inter-trial interval (ITI) of 1–2 seconds. Participants were instructed to internalize the duration without relying on counting strategies. During the first session, this phase was repeated for an additional 10 trials if the target interval had not yet been reliably memorized. In the second session, all participants completed a full set of 10 trials to ensure consistency.

Building on this internal representation, the second phase focused on time production (TP). Participants completed 30 trials in which they reproduced the target duration by using a computer mouse to mark both the onset and offset of the interval. Each trial began with a white fixation cross indicating the start of the response. Following each attempt, visual feedback informed participants whether their reproduction fell within ±1000 ms of the target; trials exceeding this threshold were classified as invalid and excluded from further analysis. This TP training phase was not repeated in the second session.

Finally, the third phase introduced metacognitive evaluation through temporal error monitoring (TEM). In 10 trials, participants first reproduced the target duration and then provided two judgements: (i) an estimate of the direction and magnitude of their timing error on a continuous scale, and (ii) a confidence rating for that estimate. Both responses were collected using a custom arc-based interface (Figure 1), in which participants dragged the mouse from the center toward a position on the arc, allowing for high-resolution and spatially unbiased input.

To facilitate calibration of these metacognitive judgements, participants received explicit feedback on the magnitude of their timing errors during this phase. Feedback was individualized based on each participant’s performance variability, quantified as the standard deviation (SD) of TP errors obtained in the second phase. Errors were categorized into four levels: *too short/long* (>2 SD), *quite short/long* (>1 SD), *a little bit short/long* (>0.3 SD), and *very close to the target* (≤0.3 SD). On Day 2, participants completed an additional 15 TEM training trials, with feedback calibrated using the TP standard deviation derived from Day 1.

#### b) Main Experimental Phase

During the main phase, each trial began with participants initiating the interval by pressing a button and terminating it with a second press once they judged that 2 seconds had elapsed, constituting the TP. Following this, they provided two metacognitive judgements. The TEM phase was separated from the TP response by a brief 300 ms interval, measured from the offset of the produced interval to the onset of the judgement phase.

Feedback availability was manipulated within the session: 50% of trials included feedback, while the remaining 50% did not. On feedback-present trials, participants received performance feedback based on the same individualized deviation categories established during TEM training. In contrast, on feedback-absent trials, no external information was provided, requiring participants to rely exclusively on their internal representations of time to evaluate their performance. This manipulation enabled a direct assessment of TEM under conditions with and without external feedback.

##### Computation of TEM variables and confidence ratings

Both judgements were recorded via the arc-based interface introduced during training. The arc comprised 101 evenly spaced points spanning -45 ° to +45 °. For each mouse click, the arc coordinate with the smallest Euclidean distance to the click position was selected as the raw response. This response was then mapped onto a standardized continuous scale. Temporal error estimates were normalized from −100% to +100%, where −100% indicated maximum underestimation and +100% maximum overestimation. Confidence ratings were normalized from 0% to 100%. This ensured both measures were represented on continuous, trial-specific scales suitable for single-trial analysis.

Behavioral metrics extracted from each trial included TP duration, temporal error response, and TEM confidence rating. A Metacognitive Distance (MD) index was computed as the absolute difference between the z-scored temporal error response and the z-scored TP duration, providing a trial-wise measure of alignment between temporal error estimates and actual timing performance. Formally, for trial (*i*):

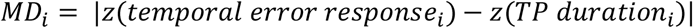

##### TP categorization

Continuous TP values were discretized into three groups using the 33rd and 66th percentiles of the TP distribution as thresholds. Trials below the lower threshold were labelled “Short”, those above the upper threshold were labelled “Long”, and trials between the two thresholds were labelled “Correct”. This tertile-based categorization yielded an approximately balanced distribution across the three classes and served as the multiclass classification target.

##### MI categorization

MD values were dichotomized using a median split across all trials. Trials below the median were classified as “High MI” and those at or above as “Low MI”, on the assumption that smaller absolute differences reflect more accurate metacognitive inference. This yielded approximately equal trial counts across groups and defined the binary classification target.

##### TEM confidence categorization

TEM confidence values were similarly dichotomized using a median split. Trials below the median were assigned to the “Low Confidence” group and those at or above to the “High Confidence” group, providing a balanced binary classification target for subsequent analyses.

Overall, these categorizations enabled evaluation of motor action timing performance and metacognitive precision at the single-trial level.

## 3) Statistical Analysis

Behavioral data were analyzed in Python (version 3.12) using the *statsmodels* package^26^ (version 0.14.5). Statistical analyses included ordinary least squares regression, analysis of variance, and pairwise comparisons. Single-trial measures of TP and TEM were preprocessed by excluding outliers exceeding 3.5 standard deviations from each participant’s mean.

Relationships among continuous variables were assessed using linear mixed-effects models (LMMs) estimated via restricted maximum likelihood (REML). Trial-level observations were nested within participants to account for repeated measures. Fixed effects captured population-level relationships, whereas random effects modeled between-participant variability. Model selection was guided by the Akaike Information Criterion (AIC) and likelihood ratio tests for nested comparisons. Model-derived slopes were visualized with 95% confidence intervals.

Group-level differences in decoding accuracy and behavioral measures were evaluated using independent-samples *t*-tests. This analytical framework enabled robust estimation of effects while appropriately accounting for repeated measures and inter-individual variability.

## 4) EEG Data Processing

### 4.1) Signal Denoising

Raw EEG data were processed using MNE-Python^28,29^ (v1.9.0). Data were loaded from BrainVision files, and irrelevant segments, such as inter-block breaks, were removed. One participant (nr. 18) was excluded due to excessive α-wave activity, yielding a final sample of 28 participants.

EEG signals were initially re-referenced to the average of all channels to eliminate global shifts. Poor-quality channels were identified and excluded before Independent Component Analysis (ICA). A 50 Hz notch filter was applied to remove power-line interference, and a bandpass filter (1–120 Hz) was applied to remove low-frequency drift and high-frequency noise. ICA decomposed the signal into independent components, which were visually inspected to identify and remove artifact-related components, including eye movements (EOG), muscle contractions (EMG), and heartbeats (ECG). Bad channels were subsequently interpolated, and data were re-referenced to the channel average.

Artifact rejection employed the AutoReject protocol^30^ (v0.4.2), with interpolation levels (1, 4, 16, 32) and consensus percentages (0–1.0) optimized via Bayesian methods. Time-frequency analysis was performed using multitaper decomposition across θ (3–7 Hz), α (8–12 Hz), and β (13–30 Hz) bands, as well as a merged band combining all three. Cycle parameters for each band were selected to balance time and frequency resolution, reducing spectral leakage and improving frequency estimation precision.

Baseline correction was applied relative to a reference window of −0.3 to −0.1 s before the first keypress. Time-frequency data were then cropped from TP onset to 1.2 s post-onset, capturing frequency dynamics during the TP phase. The TP offset marker was excluded to ensure that model predictions relied solely on ongoing EEG activity rather than the interval termination.

### 4.2) Signal Filtering for Model Input

#### a) Raw EEG Data Structure

EEG data were preprocessed and analyzed at the individual-participant level to account for inter-subject variability in noise characteristics and neural activity patterns.^31^ This participant-specific approach mitigates a well-established limitation of group-level EEG analyses, in which pooling data across participants can obscure fine-grained, trial-level neural dynamics relevant to the decoding of cognitive states.

Following denoising and artefact correction, clean EEG signals were organized into a four-dimensional tensor:

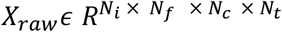

where *N*_*i*_ denotes the number of trials, *N*_*f*_ the number of frequency bins derived from time-frequency decomposition, *N*_*c*_ the number of EEG channels, and *N*_*t*_ the number of time points capturing within-trial temporal dynamics (Figure 2a). This representation preserves the joint spectral, spatial, and temporal structure of the EEG signal, which is frequently collapsed or averaged in conventional preprocessing pipelines.

**Figure 2.**
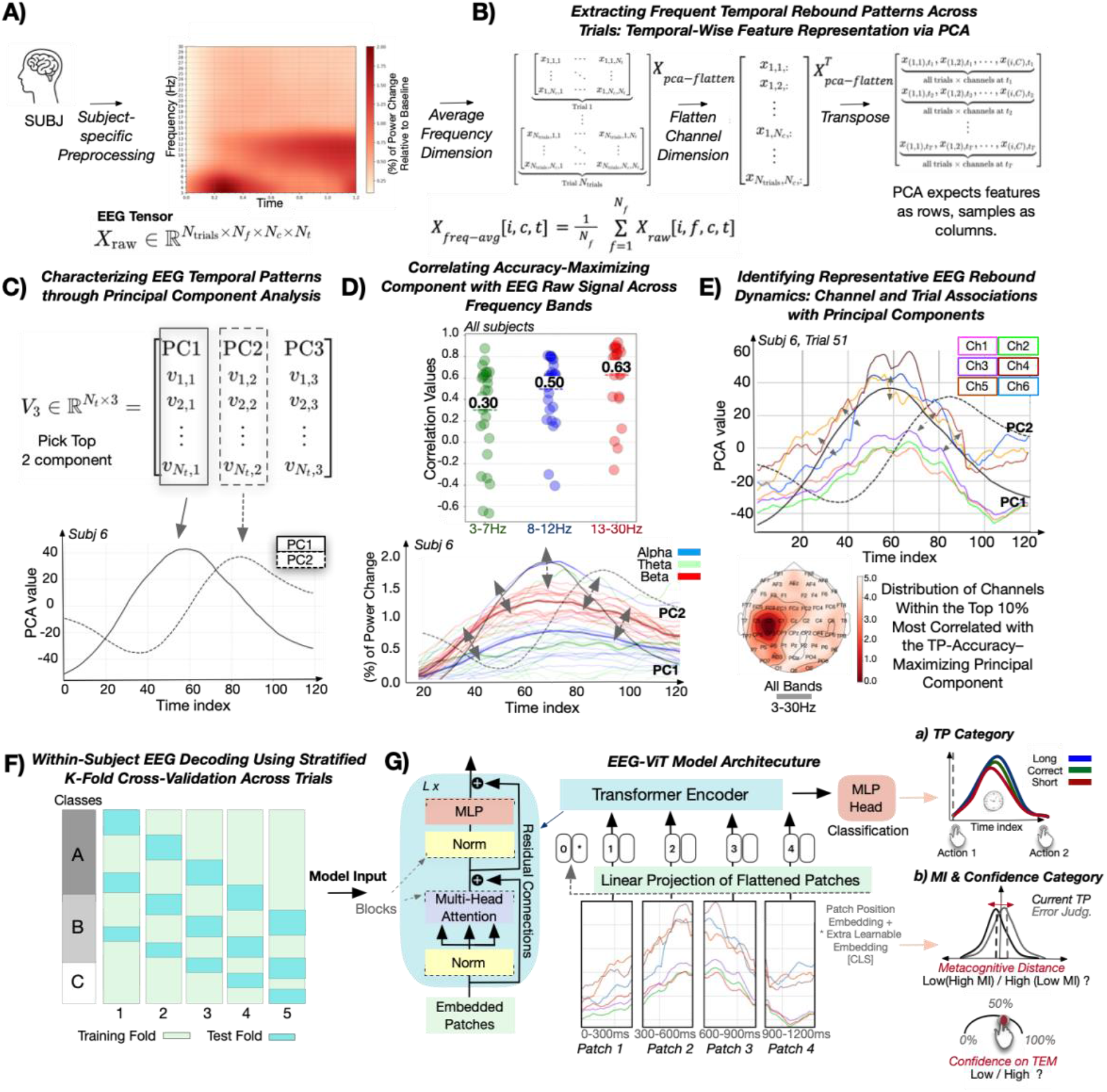
Subject-specific preprocessing pipeline and Vision Transformer architecture for EEG-based behavioral decoding. **a EEG preprocessing and data structuring**. Raw EEG underwent artifact rejection, bandpass filtering (1–120 Hz), notch filtering (50 Hz), and baseline normalization. Preprocessed signals were structured as four-dimensional tensors (trials × channels × frequencies × time). The right panel shows a time-frequency representation averaged across participants and trials, with color indicating percentage power change relative to baseline (−0.3 to −0.1 s before TP onset). See Methods 3.1–3.2. **b Data preparation for PCA**. EEG signals were frequency-averaged across three bands (θ: 4–7 Hz; α: 8–12 Hz; β: 13–30 Hz) and flattened across channels and trials to reduce dimensionality and extract dominant temporal patterns. The resulting matrix was transposed into a trials × channels × time representation suitable for principal component analysis (PCA), with each trial treated as a feature to identify common temporal patterns across channels. See Methods 3.2a–d. **c Principal component extraction**. PCA was applied to the covariance matrix of the transposed data. Eigenvalue decomposition yielded temporal components ranked by explained variance. The first two components (PC1 and PC2) were retained to capture the dominant temporal structure across trials. Example waveforms for participant 6 are shown in the bottom panel. Post-movement β activity showed considerable variability, with no standardized temporal range defining a canonical β rebound. The accuracy-maximizing component was defined as the one yielding the highest decoding performance. Similar rebound patterns were observed in the α and θ bands. See Methods 3.2e–g. **d Validation of extracted components against physiological oscillations**. To assess whether PCA-derived components reflected physiologically meaningful activity, the accuracy-maximizing component was correlated with the raw EEG signal in each frequency band. β-band activity showed the strongest associations (*M* ≈ 0.63), followed by α (*M* ≈ 0.50) and θ (*M* ≈ 0.30). Pairwise comparisons using Tukey’s HSD confirmed that β correlations were significantly higher than θ (*MD* = −0.33, *p* = .007, 95% CI [−0.58, −0.08]), while α did not differ significantly from β (*MD* = 0.14, *p* = .396) or θ (*MD* = −0.19, *p* = .172). These findings confirm that the accuracy-maximizing component most strongly captured β-rebound dynamics, with α and θ contributing more modestly. The bottom panel illustrates this component for participant 6, with frequency-specific correlations highlighted in red (β), blue (α), and green (θ). **e Selection of informative channels and trials**. Correlations between the accuracy-maximizing component and frequency-averaged EEG signals were computed to identify the most informative signals for model input. Channels were ranked first, followed by trials, with the top 10% of channels and top 30% of trials retained per participant. Example data from participant 6, trial 51, are shown in the top panel, with each waveform representing one of the six top-ranked channels. The bottom topographic plot shows channels consistently selected across participants, with clusters over central and parietal regions overlapping with known post-movement β rebound areas, supporting the functional relevance of the selection procedure. See Methods 3.2h. **f Stratified cross-validation design**. Within-participant decoding used stratified 5-fold cross-validation, maintaining proportional representation of behavioral categories (TP, MI, and TEM confidence) across folds. This minimized class-imbalance bias and prevented information leakage between training and test sets. See Methods 4d. **g Vision Transformer architecture for temporal pattern learning**. Single-trial EEG signals (trials × channels) were segmented into contiguous temporal patches, each linearly projected into fixed-dimensional embeddings. A learnable class token ([CLS]) was prepended to the patch sequence to aggregate trial-level information, with its final state serving as the classification input, following established NLP architectures such as BERT^27^. Positional encodings preserved temporal order across the sequence. Embeddings were passed through six stacked transformer encoder layers, each containing multi-head self-attention to identify informative temporal relationships, layer normalization for training stability, residual connections for gradient flow, and feedforward MLP blocks. The final [CLS] embedding was classified via MLP to predict behavioral categories (TP, MI, and TEM confidence). This architecture captured both local and global temporal dynamics, enabling identification of distributed EEG patterns that predict trial-level behaviours. See Methods 4a–c.

For TP, MI, and TEM confidence classification based on EEG activity during the TP phase, each trial comprised 120 time points spanning 0–1.2 s relative to the event marker. For MI classification based on activity during the temporal error response and TEM confidence judgement phases, each trial comprised 60 time points spanning 0–0.6 s, reflecting the shorter temporal scale of post-response evaluative processing.

#### b) Frequency-Averaged Data Representation

EEG data were averaged across the frequency dimension to isolate temporally structured neural dynamics independently of band-specific spectral contributions, producing a three-dimensional representation that preserved trial-wise spatial and temporal information (Figure 2b):

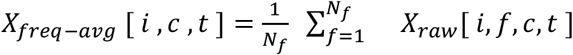

resulting in:

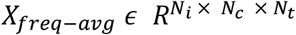

Each trial-specific slice represented the multichannel EEG time series collapsed across frequencies, with channels along rows and time points along columns. Collapsing across frequencies reduced input dimensionality and mitigated redundancy from correlated spectral components, while preserving the temporal dynamics of neural activity. This representation enabled a direct test of whether temporally organized activity alone was sufficient to decode task-relevant cognitive and metacognitive states. Model performance on frequency-resolved versus frequency-averaged inputs was then compared to assess the relative contribution of oscillatory specificity beyond broadband temporal structure. This contrast clarified the extent to which trial-by-trial variability in motor timing, error monitoring, and TEM confidence was driven by temporal patterns rather than frequency-specific neural dynamics.

#### c) Flattened Trial-Channel Matrix for PCA

To characterize dominant temporal modes of variability shared across trials and recording sites, the frequency-averaged EEG data were reshaped into a two-dimensional matrix suitable for PCA. The trial and channel dimensions were collapsed into a single observation axis:

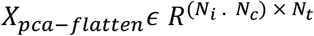

where each row corresponded to an individual trial–channel observation and each column to a time point (Figure 2b). Treating each trial–channel pair as a distinct realization of the underlying temporal dynamics preserved the full temporal structure along the feature axis. This formulation directed PCA toward low-dimensional temporal components capturing shared variance across trials and channels, rather than idiosyncratic spatial or trial-specific fluctuations.

#### d) Transposed Matrix for PCA

To extract shared temporal structure, the flattened matrix was transposed such that time points served as variables and trial–channel pairs as observations:

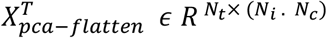

Each row thus represented the percentage change in EEG power relative to baseline across all trial–channel observations at a given time point (Figure 2b). This orientation directed PCA towards capturing temporal correlations across trials and channels, which enabled the identification of common temporal patterns.

#### e) Covariance Matrix

After mean-centering the transposed matrix across observations,

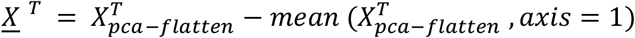

the temporal covariance matrix was computed as:

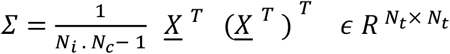

Each element 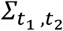 quantified the covariance between time points *t*_1_ and *t*_2_ across all trial–channel observations. Diagonal elements reflected temporal variance, while off-diagonal elements captured temporal correlations. This covariance structure formed the basis for extracting principal temporal components.

#### f) Principal Components

Eigen-decomposition of the covariance matrix ∑ yielded eigenvectors *V* and eigenvalues *Λ*:

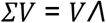

where 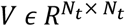 contained eigenvectors as columns and *Λ* was the corresponding diagonal eigenvalue matrix. The first three eigenvectors,

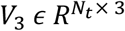

represented the dominant temporal activity patterns, with each column corresponding to a principal component (PC1, PC2, PC3) and each row to a time point (Figure 2c). These components summarized the primary temporal structure shared across trials and channels, accounting for the largest proportion of total variance.

#### g) PCA Scores

Each trial–channel observation was projected onto the dominant temporal components by multiplying the flattened matrix by the top eigenvectors:

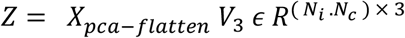

Each row *Z* represented a trial–channel observation, and each column its loading on one of the three principal temporal components. This produced a compact representation of the EEG data while retaining both trial- and channel-level information.

#### h) Channel and Trial Selection Based on Principal Components’ Correlations

EEG trials and channels vary in how strongly they express dominant temporal patterns, reflecting differences in neural responses, cognitive state, and noise. The first two principal components (PC1 and PC2, collectively *V*_2_) were extracted from the frequency-averaged EEG data to isolate the most informative and reproducible signals. PC1 and PC2 captured dominant temporal dynamics consistent with β-band rebound activity, previously implicated in motor action timing^17,18^ and temporal error monitoring.^4^

For each component *V*_*k*_ (*k* = 1, 2), the average correlation between the component and each channel’s EEG signal across all trials was computed:

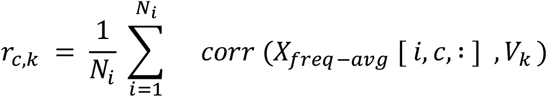

Channels were ranked by their average correlations with PC1 and PC2, and the top 10% (six channels) were retained for further analysis (Figure 2e):

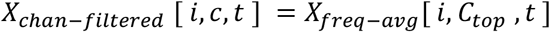

For each trial *i*, the retained channel signals were averaged:

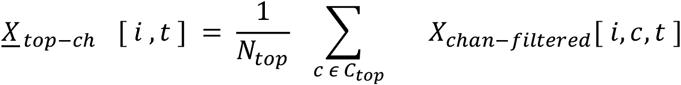

Trials were subsequently ranked by their correlation with PC1 and PC2:

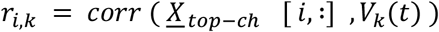

This ranking prioritized trials most strongly expressing the shared temporal dynamics of interest, reducing the influence of noisy or weakly aligned observations. Trial subsets corresponding to the top 30%, 50%, and 100% were retained to evaluate how alignment with dominant temporal patterns affects downstream model performance. The resulting trial– channel–time representations, restricted to selected channels and ranked trial subsets,

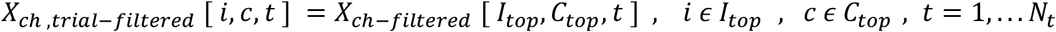

served as inputs for all subsequent predictive analyses. Comparing model performance across trial subsets showed that prioritizing channels and trials aligned with shared temporal dynamics improved decoding accuracy and the generalizability of learned neural representations.

## 5) EEG Vision Transformer (EEG-ViT) Model Pipeline

### a) Model Input Representation

Following channel and trial selection, the preprocessed EEG data were organized as:

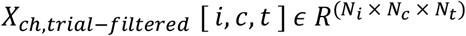

This filtered dataset retained only the most informative trials and channels, improving signal-to-noise ratio and focusing downstream modelling on dominant temporal patterns. Each trial– channel pair was treated as an independent observation, and the 3D tensor was reshaped into a 2D matrix:

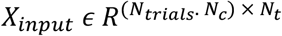

Each row corresponded to the temporal EEG signal from a single trial–channel pair, and each column to a time point. This reshaping emphasized temporal structure while allowing patterns to be learned jointly across trials and channels, without imposing a fixed spatial topology.

### b) Patch Embedding and Input Construction

Transient, band-limited power rebounds, such as β-band rebounds, are hallmark neural signatures of motor action timing and temporal error monitoring. These events are brief, variable in latency and amplitude across trials and channels, and embedded within substantial background noise. Their reliable detection requires two complementary capacities: *(i)* local representations capturing short temporal segments of power modulation, and *(ii)* a global integration mechanism that combines local signals across the trial to form a coherent representation of the underlying neural state.

To satisfy both requirements, the one-dimensional EEG time series was treated as *a temporal image* and modelled using a Vision Transformer (ViT) architecture adapted for single-trial EEG,^22^ with our minor modifications to accommodate trial-specific temporal dynamics (Figure 2g). Temporal patch segmentation is conceptually analogous to spatial patch decomposition in image ViTs: each patch encodes local temporal (and implicitly spectral) structure within a short window, while multi-head self-attention captures interactions between patches across the full trial. Critically, unlike convolutional filters, which impose fixed local receptive fields, self-attention can model dependencies across arbitrary temporal distances, enabling the detection of events whose diagnostic relevance emerges from broader temporal context (e.g., a rebound following earlier suppression).

The ViT was configured with a grid size of (1,1) and a spatial patch size of (1,1), treating each channel as a single spatial unit. Each temporal sequence was divided into *N* = *T*/*P*_*t*_ = 4 non-overlapping patches of length *P*_*t*_ = 30 time points (see Table 1 for full hyperparameters). For sample *i*, patch *j* was defined as:

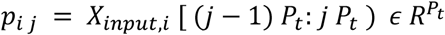

**Table 1.** Hyperparameters for the ViT-based EEG decoding pipeline. The table depicts the hyperparameters and selected values used in the ViT-based EEG decoding pipeline for patch embedding, transformer architecture, and model training, together with brief notes on parameter ranges.

| Parameter | TPOnsetStart Event | TEMOnsetStart Event |
| --- | --- | --- |
| Input chunk size | 120 time points | 60 time points |
| Temporal patch size (Pt) | 30 | 15 |
| Number of patches (N) | 4 | 4 |
| Embedding dimension (d) | 64 | 64 |
| Transformer depth (L) | 6 layers | 6 layers |
| Attention heads (H) | 6 | 6 |
| Head dimension (dh) | 32 | 32 |
| Feed-forward channels (dff) | 64 | 64 |
| Dropout rate | 0.15 | 0.15 |
| Embedding dropout | 0.15 | 0.15 |
| Grid size | (1, 1) | (1, 1) |
| Spatial patch size | (1, 1) | (1, 1) |
| Output classes | TP (Short, Correct, Long); MI (Low, High); Confidence (Low, High) | MI (Low, High); Confidence (Low, High) |
| Epochs | 120 | 120 |
| Batch size | 18 | 18 |
| Optimizer | AdamW (lr=5e-4, weight_decay=1e-2) | AdamW (lr=5e-4, weight_decay=1e-2) |
| Learning rate scheduler | CosineAnnealingLR (Tmax=50, eta_min=1e-3) | CosineAnnealingLR (Tmax=50, eta_min=1e-3) |

Each patch was linearly projected into a 64-dimensional embedding space (*d* = 64) via a learnable mapping:

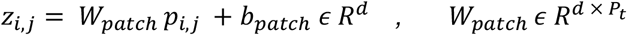

A learnable classification token [*CLS*] ϵ *R*^*d*^ was prepended to the patch sequence, and learnable positional encodings *pos* ϵ *R*^(*N*+1) × *d*^ were added to preserve temporal order:

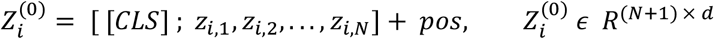

Dropout (*p* = 0.15) was applied to the embedded sequence to improve regularization.

### c) Transformer Encoder

The transformer encoder comprised *L*= 6 identical layers, each consisting of a multi-head self-attention (MHA) block followed by a position-wise feed-forward network (MLP), with residual connections and layer normalisation applied at each stage.

#### Multi-head Self-Attention (MHA)

For a layer *l*, the input *X*^(*l*−1)^ϵ *R*^*B* × (*N*+1) × *d*^ was normalized:

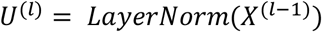

For each head *h*, queries (*Q*), keys (*K*), and values (*V*) were obtained via linear projections:

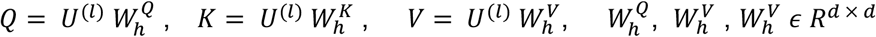

Attention was computed independently across *H* = 6 heads, each with a per-head dimension *d*_*h*_ = 32:

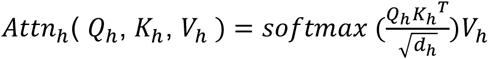

Head outputs (*W*^*o*^) were concatenated and projected:

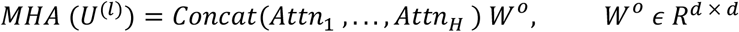

with residual update:

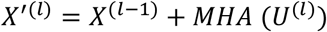

#### Feed-Forward (MLP) Block

The normalized intermediate representation,

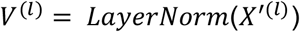

was passed through a position-wise MLP with a hidden dimension *d*_*ff*_ = 64:

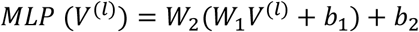

with residual update:

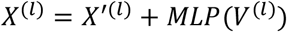

This process was repeated for *l* = 1, . . ., *L*.

#### Pooling and Classification

Following the final encoder layer, the classification token embedding was extracted:

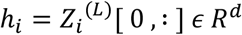

This vector provided a compact summary of the temporally distributed EEG dynamics for each trial–channel pair and was passed to a linear classifier to produce task-specific predictions:

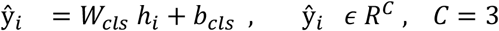

corresponding to the target label categories (e.g., short, correct, long for TP; low versus high for TEM confidence) (Figure 2g, insets a–c).

### d) Model Fitting

A stratified k-fold cross-validation (k = 5) was applied at the trial level to ensure balanced representation of both dataset size and target categories across folds (Figure 2f). In each iteration, k−1 folds were designated for training, and the remaining fold for testing, with the model reinitialized before each fold to prevent leakage of optimized parameters. Constructing folds at the trial level ensured that no trial in the test set appeared in the training set, eliminating artificial inflation of performance due to correlations across channels within the same trial. Stratification further mitigated class imbalance, yielding stable and unbiased evaluation metrics^32^.

Batches lacking examples from all target categories were skipped to prevent gradient updates from favoring overrepresented classes. This cross-validation framework maximized data utilization, balanced the bias–variance trade-off, and produced reproducible estimates of model performance across trials and participants. Evaluating accuracy across independent folds confirmed the generalizability of temporally structured neural representations, supporting applicability to diverse datasets, experimental paradigms, and cognitive processes^33^.

## 6) Results

### 6.1) Decoding Time Production and Metacognitive Processes from EEG Using ViT-Based Modeling

#### Influence of Trial Selection on TP Decoding

We first examined whether selectively training the model on trials whose oscillatory dynamics most closely matched the dominant neural pattern would improve TP category decoding. Motor action timing engages rebound dynamics across β, α, and θ frequency bands; however, these signals fluctuate substantially across trials in latency, amplitude, and spatial distribution, meaning that not all trials carry equally reliable behaviorally relevant information. The accuracy-maximizing principal component, defined as the component yielding the highest decoding performance, captured this dominant oscillatory structure across all three bands, with β showing the strongest expression. This is consistent with the well-documented variability of post-movement β rebound timing, with mean onset around 230 ms and peak activity between 500–1000 ms after movement termination^34,35^, and single-trial latency spans of ±400 ms within individuals.^36^ Analogous rebound patterns were observed in the α and θ bands, though less pronounced, confirming that the component reflected a multi-band rather than purely β-driven signal. Trials were then ranked by their alignment with this component, and decoding was performed separately on the top 30%, top 50%, and full 100% of trials, to test whether prioritizing trials that most strongly expressed this shared oscillatory structure would yield more reliable TP category classification.

Scatter-density plots (Figure 3a) display mean 5-fold cross-validated test accuracy per participant across trial selection conditions. One-sample *t*-tests confirmed that decoding accuracy significantly exceeded chance (33%) in both the top 30% subset (*M* = .377, 95% CI [.359, .394], *p* < .001) and the top 50% subset (*M* = .354, 95% CI [.341, .367], *p* < .001). Accuracy in the 100% condition (*M* = .339, 95% CI [.328, .349], *p* = .092) was numerically above chance but did not reach statistical significance. Independent two-sample *t*-tests revealed that the top 30% subset significantly outperformed both the top 50% (*t*(28) = −2.092, *p* = .041) and 100% subsets (*t*(28) = −3.779, *p* < .001), while the difference between the top 50% and 100% subsets remained marginal (*t*(28) = −1.860, *p* = .069). Trials most strongly correlated with the accuracy-maximizing component displayed clearer β-rebound-like temporal dynamics (Figure 3a, lower panel), suggesting that selective trial inclusion enriches the neural signal available for decoding. When all frequency bands were combined, decoding accuracy reached up to 57%, subject to cross-fold variability (Figure 3b). Centro frontal, central, parietal, and occipital channels were most consistently selected under the accuracy-maximizing component criterion (Figure 3c), in line with topographies typically associated with beta-rebound activity.^4,17,18^

**Figure 3.**
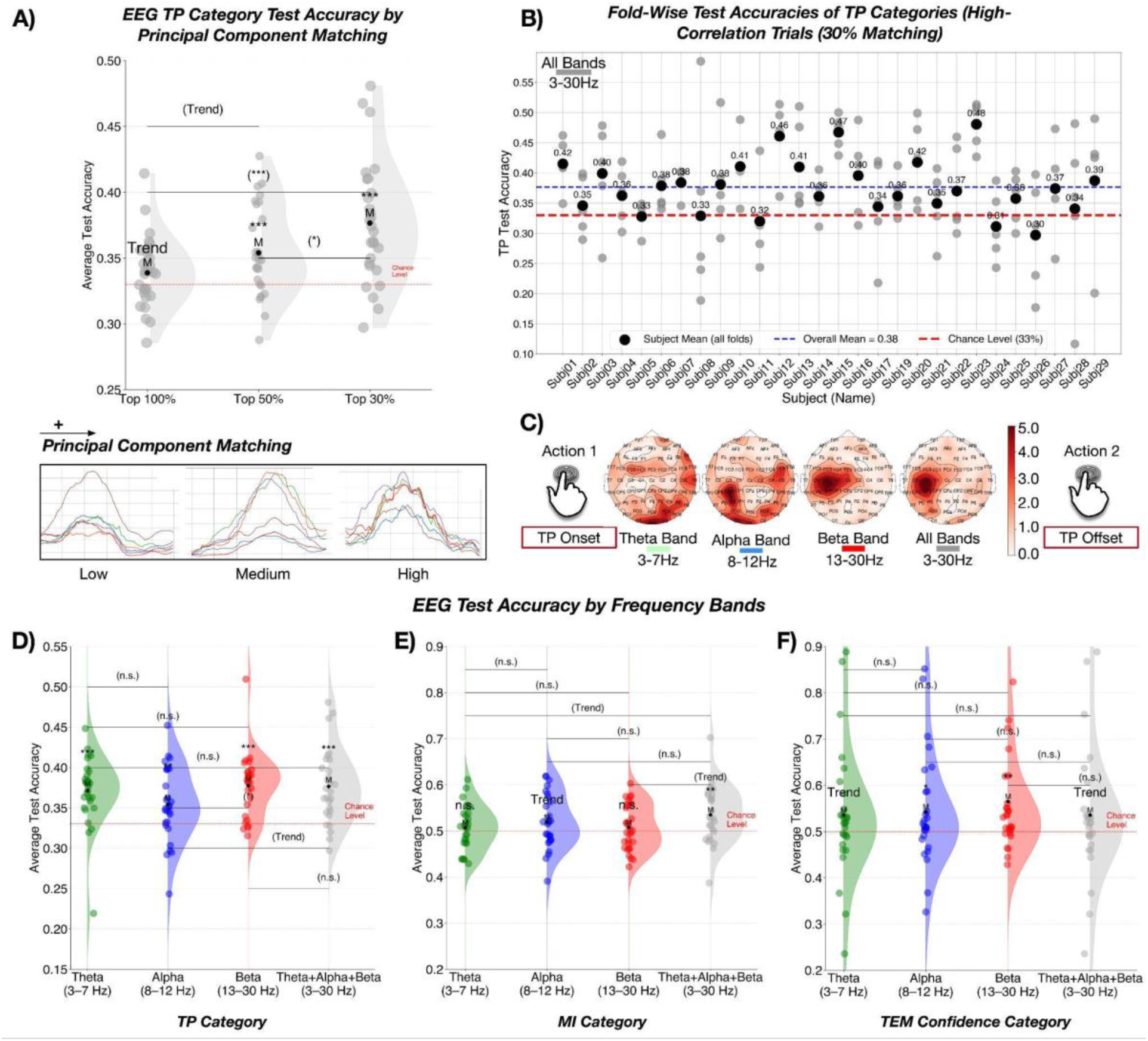
Multifrequency EEG Decoding Reveals Predictive Signals of Trial-to-Trial Time Production (TP), Metacognitive Inference (MI), and Confidence with EEG-ViT. **a Trial selection based on component alignment improves decoding accuracy**. Scatter-density plots show average 5-fold test accuracy per participant for models trained on trials ranked by correlation with the accuracy-maximizing principal component (top 30%, 50%, or 100%). P-values indicate pairwise comparisons between trial selection groups (independent two-sample t-tests). Classification accuracy increased with stricter trial selection, with the top 30% yielding the highest performance. One-sample t-tests showed that accuracy in the top 30% and top 50% exceeded chance level (33%), whereas the 100% condition showed only trend-level effects. Accuracy differences were largest between the top 30% and 100%, followed by 30% and 50%, and 50% and 100%. The lower panel shows that trials with stronger component correlations exhibited clearer temporal rebound patterns, emphasizing the importance of selective trial inclusion. **b TP categories are decodable on a single-trial basis before interval completion**. Test accuracies for TP category classification (short, accurate, long) are shown per participant using the top 30% of matched trials across all frequency bands. Colored points represent individual cross-validation fold results; black points denote fold-averaged accuracy per participant. Despite fold-to-fold variability, accuracies consistently exceeded chance level (33%; red line). The blue line indicates the grand mean across participants, confirming reliable trial-by-trial prediction of TP category before interval termination. **c Predictive neural signals exhibit frequency-dependent spatial topographies**. Topographic maps display the spatial distribution of selected channels under the top 30% matching criterion for theta (θ; 3–7 Hz; green), alpha (α; 8–12 Hz; blue), beta (β; 13–30 Hz; red), and the broadband combination (θ+α+β; 3–30 Hz; grey). β-band channel selections clustered over central and parietal regions, consistent with the established topography of post-movement β rebound. All decoding relied exclusively on neural activity recorded after TP onset and before offset, with predictive information preserved within a 1.2 s post-onset window, indicating that behaviorally relevant signals emerged well before interval completion. **d Multiple frequency bands contribute independently to TP category decoding**. Density plots summarize TP category classification accuracies across frequency bands under the top 30% matching condition. Decoding accuracy significantly exceeded chance (33%) for all bands (θ, α, and β), demonstrating that TP prediction was not exclusively dependent on β-rebound activity but also reflected temporally structured oscillations in the θ and α ranges. α-band activity showed the weakest decoding performance, consistent with prior evidence linking α dynamics to general timing variability. **e MI requires cross-frequency integration**. Single-trial MI accuracy (alignment between subjective temporal error estimates and actual TP performance) was decoded before interval termination. Significant decoding was observed only when all frequency bands (θ+α+β) were combined, with α contributing at the trend level. Individual frequency bands alone did not support reliable decoding. These findings indicate that metacognitive inference depends on the integration of information across multiple oscillatory sources, rather than being localized to a single frequency band. **f Confidence monitoring engages beta and alpha oscillatory mechanisms**. Confidence ratings associated with error monitoring were decoded before TP termination. Both α and β signals carried predictive information, with β showing the stronger effect. θ decoding was present only at the trend level. These findings suggest that confidence monitoring engages oscillatory mechanisms linked to motor-action monitoring (β) and attentional control (α).

Taken together, these findings indicate that decoding performance benefits from selectively prioritizing trials that strongly express the dominant temporal pattern of the accuracy-maximizing component. Such high-matching trials exhibit a characteristic spatiotemporal signature consistent with β-rebound dynamics, which enhances the extraction of behaviorally relevant neurophysiological features and improves TP category decoding.

#### Contributions of Frequency Bands to TP Category Decoding

Classification accuracy was compared across frequency bands under the top 30% matching condition (Figure 3d). One-sample *t*-tests confirmed significant above-chance decoding for all frequency bands: θ (*t*(28) = 5.188, *p* < .001, *M* = .371, 95% CI [.355, .388]), α (*t*(28) = 2.811, *p* = .009, *M* = .354, 95% CI [.337, .372]), β (*t*(28) = 6.439, *p* < .001, *M* = .377, 95% CI [.362, .393]), and the broadband combination θ+α+β (*t*(28) = 5.401, *p* < .001, *M* = .377, 95% CI [.359, .394]). A significant difference between α and β (*t*(28) = −2.044, *p* = .046) indicated that β-band activity carried stronger predictive information for TP categorization. A trend-level difference was observed between α and θ+α+β (*t*(28) = −1.829, *p* = .073). No other pairwise comparisons reached significance, suggesting that broadband combination did not substantially improve decoding accuracy beyond β alone.

#### Decoding MI categories

Having established that TP categories were reliably decodable from individual frequency bands, we next examined whether the same approach could decode metacognitive inference (MI), the trial-by-trial alignment between subjective error reports and objective timing performance (Figure 3e). This distinction is theoretically critical: single-process accounts predict MI should be decodable from the same bands as TP, given shared decision signals, whereas HOR theory predicts that metacognitive inference requires integration across multiple oscillatory sources. MI categories were therefore decoded using the same ViT architecture across θ, α, and β bands individually and in combination, enabling a direct test of whether first- and second-order neural representations draw on the same or distinct frequency-band resources.

Decoding results revealed a clear dissociation from the TP pattern. Significant above-chance classification was observed exclusively for the broadband combination (θ+α+β; *M* = .535, *t*(28) = 2.976, *p* = .006, 95% CI [.511, .559]), while no individual band supported reliable decoding: α (*M* = .519, *t*(28) = 1.754, *p* = .091), β (*M* = .508, *t*(28) = 0.886, *p* = .383), and θ (*M* = .507, *t*(28) = 0.786, *p* = .439). Trend-level differences were observed between the broadband combination and both θ (*t*(28) = −1.891, *p* = .064) and β (*t*(28) = −1.888, *p* = .064) individually. These results indicate that reliable MI decoding requires simultaneous integration across all three oscillatory sources, a fundamentally different pattern from TP decoding, where individual bands were each sufficient. Consistent with this, the data-driven principal component features captured trial-to-trial stability of each band’s spatiotemporal pattern with respect to TP prediction, with β contributing most strongly, followed by α and θ (Figure 3d).

#### Decoding Confidence Judgments

Having dissociated the frequency-band requirements for TP and MI decoding, we next examined whether confidence judgments, participants’ subjective certainty about their own error estimates, could be decoded from pre-termination oscillatory activity. Confidence represents a distinct metacognitive signal from MI: whereas MI captures the accuracy of error evaluation, confidence reflects the subjective sense of certainty accompanying it. Decoding confidence from the same pre-termination EEG, therefore, allowed us to ask whether subjective certainty draws on the same oscillatory resources as metacognitive inference, or whether it is supported by a partially separable neural signature.

Confidence ratings deviated significantly from normality across all frequency bands (Shapiro-Wilk: θ, *p* = .010; α, *p* = .010; β, *p* < .001; θ+α+β, *p* = .010) and were therefore analyzed using nonparametric tests (Figure 3f). Wilcoxon signed-rank tests indicated significant above-chance decoding for β (*W* = 328, *p* = .002) and α (*W* = 284, *p* = .033), while θ (*W* = 266, *p* = .078) and the broadband combination (*W* = 266, *p* = .078) reached only trend level. Mann-Whitney U tests revealed no significant differences in decoding accuracy across bands. Unlike MI, which required simultaneous integration across all three bands, confidence was primarily supported by α- and β-band dynamics, with β showing the stronger effect, consistent with prior evidence linking β-band activity to post-movement evaluation and internal performance monitoring. This pattern suggests that subjective certainty draws on a partially distinct oscillatory substrate from metacognitive inference itself.

### 6.2) Neural predictability of metacognitive states maps onto behavioral error monitoring

Two competing theoretical accounts generate opposing predictions about the relationship between pre-termination neural activity and subsequent error monitoring behavior, yet direct neural evidence adjudicating between them has been lacking. Single-process accounts propose that first-order performance and metacognitive evaluation arise from a shared decision variable, predicting that participants whose pre-termination neural activity more accurately encodes timing performance should also exhibit stronger error monitoring.^12,16^ HOR theory, by contrast, predicts a selective dissociation: no such relationship for TP decoding accuracy, but a specific link between MI decoding accuracy and error monitoring strength.^14,15^ To provide this first direct test, we examined whether inter-individual differences in pre-termination neural decoding accuracy, separately for TP and MI categories, predicted subsequent behavioral error monitoring (Figure 4a). Two decoding targets were distinguished: TP category decoding accuracy, reflecting the extent to which pre-termination neural activity predicted first-order timing outcomes, and MI category decoding accuracy, reflecting the extent to which it predicted the trial-by-trial alignment between objective performance and subjective error reports.

**Figure 4.**
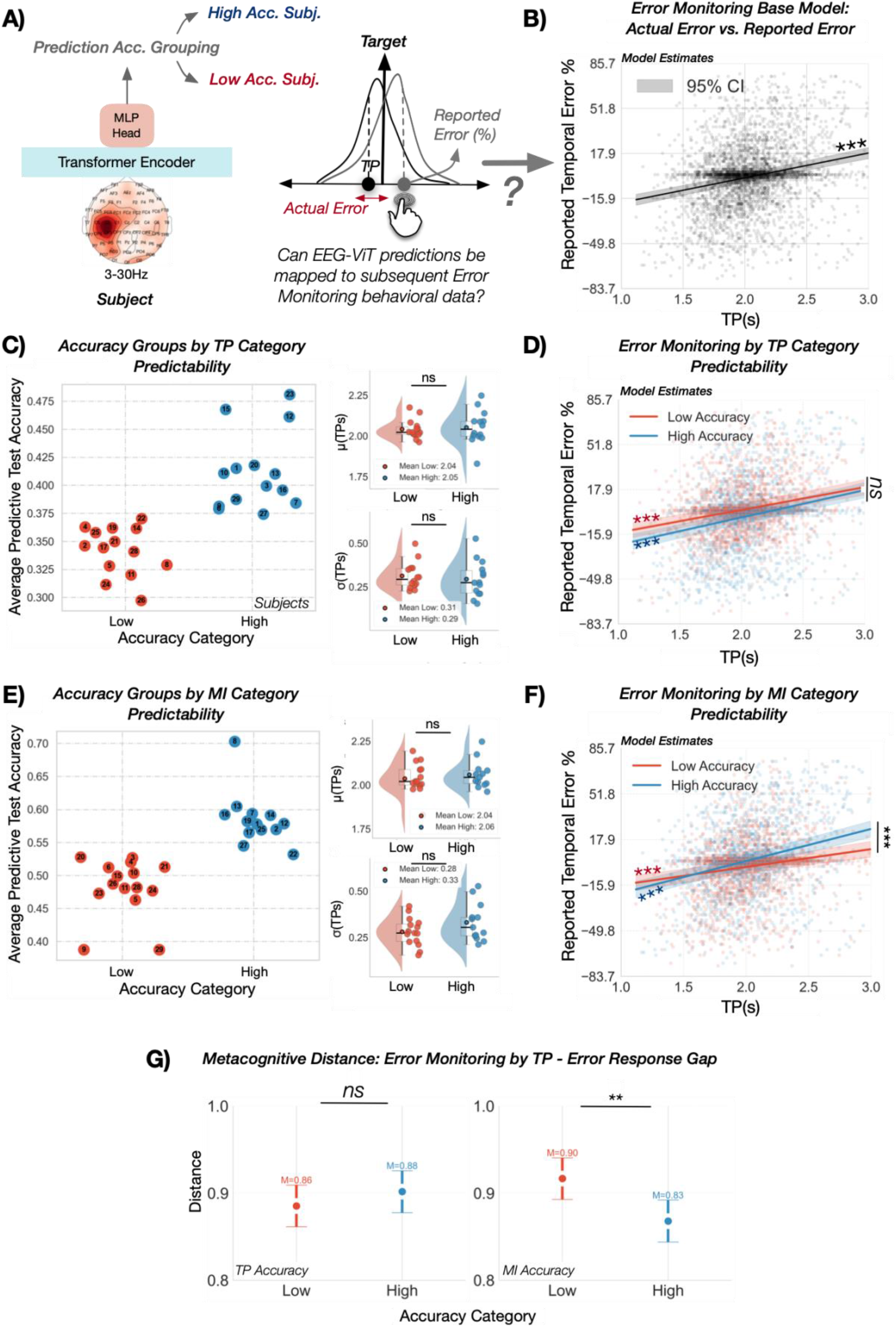
Pre-Termination Neural Decoding of Metacognitive States Predicts Behavioral Error Monitoring. **a Decoding-based stratification pipeline linking pre-termination neural dynamics to behavioral error monitoring**. Pre-termination refers to neural activity recorded during ongoing motor action timing, before the button press that terminates the produced interval. This activity was decoded using EEG-ViT to classify TP and MI categories. Reported error reflects participants’ subjective temporal error estimates converted to percentages, where sign indicates deviation direction (negative = shorter than target; positive = longer) and magnitude quantifies error size. MI category accuracy captures the trial-by-trial alignment between objective timing performance and subjective error reports. Participants were stratified into High and Low decoding accuracy groups based on model classification performance for each target separately, enabling independent tests of two competing predictions: single-process accounts predict that stronger TP neural decoding should correlate with stronger behavioral error monitoring, whereas HOR theory predicts no such relationship for TP decoding but a selective one for MI decoding. **b Trial-level behavioral validation: temporal deviations predict reported error**. The relationship between TPs and reported error percentage was examined at the single-trial level using linear mixed-effects models. The solid black line shows the population-level fixed-effect estimate, and the shaded area indicates the 95% confidence interval. Random intercepts modelled participant-level variability across repeated measurements. Scatter points represent individual trial data from all participants. The significant positive correlation indicates that deviations in either direction (shorter or longer TPs) were associated with higher reported error percentages, confirming effective trial-by-trial error monitoring. Statistical significance: ***p < 0.001, **p < 0.01, *p < 0.05, ns p > 0.05. **c Participant stratification by TP category decoding accuracy**. The strip plot shows individual participants’ average pre-termination EEG-ViT classification accuracy for TP categories, stratified into Low (red) and High (blue) TP decoding accuracy groups. Each dot represents one participant, with horizontal jitter applied for visibility and subject identifiers displayed within markers. Insets show group-level timing-response means (top) and standard deviations (bottom), revealing no significant differences in behavioral performance between the Low and High TP decoding accuracy groups. **d TP category decoding accuracy does not modulate error monitoring strength**. Model-implied correlations between TPs and reported error percentage are shown separately for the Low and High TP decoding accuracy groups. Scatter points represent individual trial data for the High (blue) and Low (red) decoding accuracy groups. Both groups exhibited positive correlations with comparable slopes. Contrary to single-process predictions, TP decoding accuracy did not significantly correlate with differences in subsequent behavioral error monitoring strength. **e Participant stratification by MI category decoding accuracy**. The strip plot shows individual participants’ average pre-termination EEG-ViT classification accuracy for MI categories, stratified into Low (red) and High (blue) MI decoding accuracy groups. Visualization conventions follow panel c. Insets display group-level timing performance means (top) and standard deviations (bottom), indicating no significant differences in behavioral performance between the Low and High MI decoding accuracy groups. **f Higher MI category decoding accuracy selectively predicts stronger behavioral error monitoring**. Model-implied correlations between TPs and reported error percentage are shown separately for the Low and High MI decoding accuracy groups. Both groups showed positive correlations, but the correlation was significantly stronger in the High MI decoding group. Consistent with the HOR theory, participants whose pre-termination neural dynamics more reliably decoded MI categories exhibited stronger behavioral error monitoring. This establishes a direct link between neural decoding of metacognitive states and subsequent error monitoring behavior. **g Metacognitive distance is reduced in the high MI category decoding accuracy group**. Bar plots show metacognitive distance (MD), computed as the difference between z-scored temporal error responses and z-scored TPs, providing a trial-wise measure of alignment between participants’ error monitoring estimates and actual motor timing performance. Error bars indicate the 95% confidence interval of the group mean. MI categories were dichotomized using a median split of MD values across all trials: trials below the median were classified as High MI (more accurate metacognitive inference) and trials at or above the median as Low MI, on the assumption that smaller differences reflect more accurate metacognitive inference. MD was significantly lower in the High MI decoding accuracy group than in the Low MI group, but only when participants were stratified by MI decoding accuracy and not by TP decoding accuracy. Together, these findings support the higher-order representation (HOR) account: more accurate pre-termination neural decoding of metacognitive states predicted superior subsequent metacognitive performance, reflected in a closer alignment between objective performance and reported error percentage.

Before testing these theoretical predictions, we confirmed reliable trial-level behavioral error monitoring. Produced intervals were significantly correlated with reported error percentage (*b* = 18.69, *SE* = 1.25, *z* = 14.97, *p* < .001, 95% CI [16.24, 21.13]; Figure 4b), confirming that subjective error evaluations systematically tracked actual performance on a trial-by-trial basis.

To test single-process predictions, participants were stratified into Low and High TP decoding accuracy groups (Figure 4c), which did not differ in mean timing accuracy (*t* = −0.34, *p* = .737) or variability (*t* = 0.54, *p* = .593). Despite robust inter-individual differences in TP decoding accuracy, the correlation between produced intervals and reported error was comparable across groups, with no significant interaction (*b* = 3.54, *SE* = 2.51, *z* = 1.41, *p* = .158, 95% CI [−1.37, 8.46]; Figure 4d). Including decoding accuracy group as an interaction term did not improve model fit (ΔAIC = −0.02; χ^2^(1) = 2.02, *p* = .155), inconsistent with single-process predictions.

MI decoding accuracy, by contrast, was selectively associated with behavioral error monitoring, consistent with HOR theory. Participants stratified into Low and High MI decoding accuracy groups (Figure 4e) did not differ in mean timing accuracy (*t* = −0.70, *p* = .488) or variability (*t* = −1.52, *p* = .141), ruling out performance-level confounds. Although both groups showed positive correlations between produced intervals and reported error, this relationship was significantly stronger in the High MI decoding accuracy group (*b* = 10.61, *SE* = 2.51, *z* = 4.24, *p* < .001, 95% CI [5.70, 15.52]; Figure 4f), with the inclusion of decoding accuracy group as an interaction term substantially improving model fit (ΔAIC = −15.83; χ^2^(1) = 17.83, *p* < .001).

This dissociation was further validated using metacognitive distance (MD), defined as the absolute difference between z-scored reported error and z-scored produced intervals (Figure 4g). Stratification by TP decoding accuracy yielded no group difference in MD (*t* = −0.949, *p* = .343; Low: *M* = 0.857, 95% CI [0.827, 0.887]; High: *M* = 0.877, 95% CI [0.847, 0.907]). Stratification by MI decoding accuracy, however, revealed significantly lower MD in the High group (*t* = 2.809, *p* = .005; Low: *M* = 0.896, 95% CI [0.866, 0.926]; High: *M* = 0.835, 95% CI [0.805, 0.865]), reflecting superior alignment between objective performance and subjective error estimates.

Together, these findings reveal a functional dissociation in the behavioral relevance of pre-termination oscillatory activity: TP decoding accuracy did not modulate subsequent error evaluation, whereas MI decoding accuracy predicted both stronger performance–error correlations and reduced metacognitive distance (Figures 4f and 4g).

### 6.3) Metacognitive inference was strongly decoded from distributed EEG activity during the TEM decision phase

The preceding analyses established that pre-termination oscillatory dynamics selectively predicted behavioral error monitoring. A remaining question is whether metacognitive signals were specific to the action execution phase or whether they strengthened further as participants transitioned into explicit error evaluation. To address this, we compared MI decoding accuracy between the TP onset and TEM decision phases across all frequency bands, allowing us to characterize both the temporal evolution of metacognitive neural signals and the consistency of their frequency-band requirements across task stages.

Classification accuracy during the TEM decision phase demonstrated that MI could be reliably decoded from all frequency bands: *θ* (*t*(28) = 6.548, *p* < .001, *M* = 0.556, 95% CI [0.538, 0.573]), *α* (*t*(28) = 6.763, *p* < .001, *M* = 0.552, 95% CI [0.536, 0.567]), *β* (*t*(28) = 6.398, *p* < .001, *M* = 0.544, 95% CI [0.530, 0.558]), and the broadband combination θ+α+β (*t*(28) = 8.434, *p* < .001, *M* = 0.575, 95% CI [0.557, 0.593]; Figure 5b). Combined-band decoding significantly exceeded both *α* alone (*t*(28) = −1.980, *p* < .05) and *β* alone (*t*(28) = −2.713, *p* < .01), but not *θ* alone (*t*(28) = −1.559, *p* = .091), suggesting that *θ*-band activity plays a particularly prominent role in metacognitive processing and that cross-frequency integration provides maximal predictive information.

**Figure 5.**
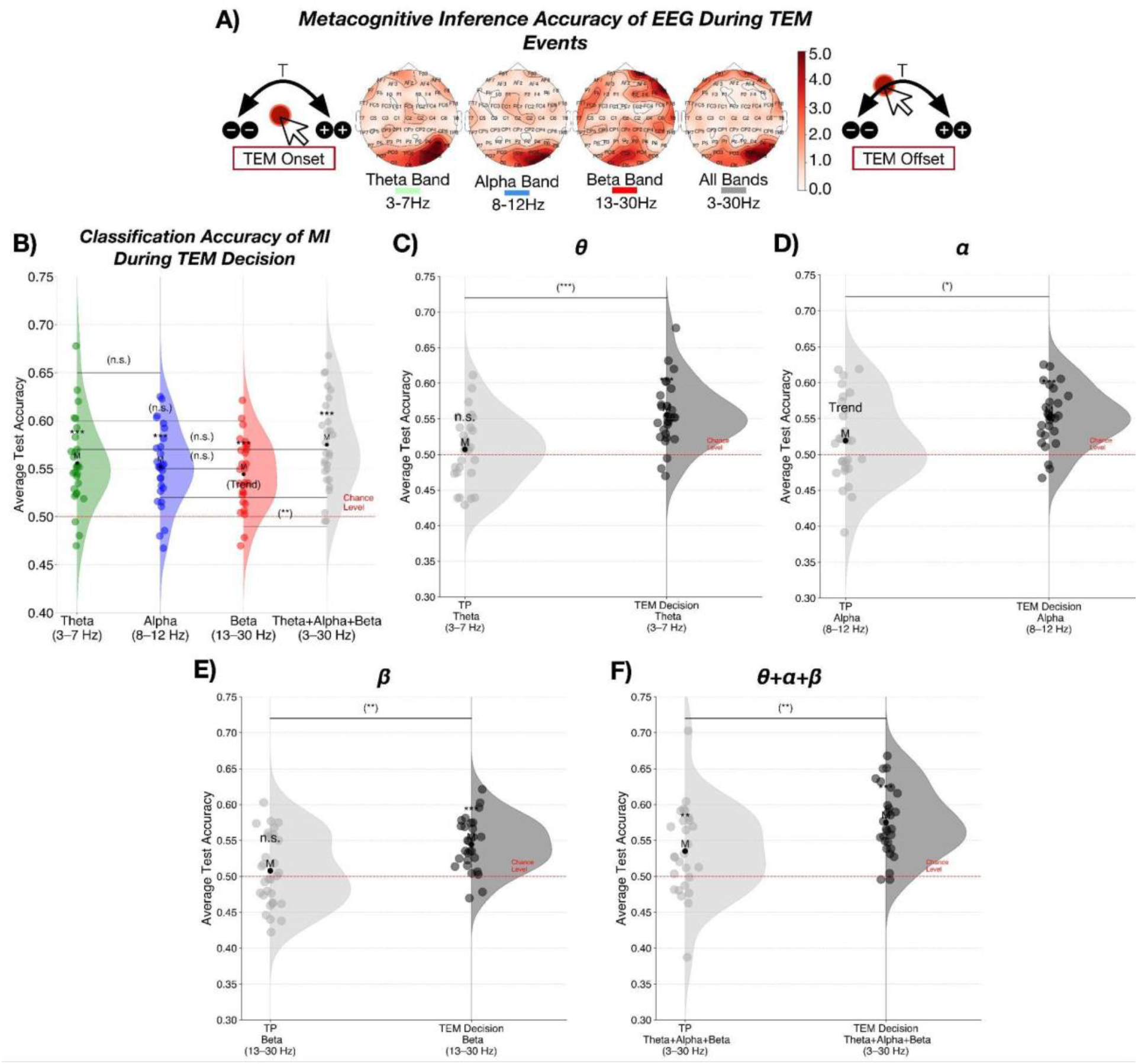
Distributed Oscillatory Signals Track MI During the TEM Decision Phase. **a Parieto-frontal topography of MI category decoding during error evaluation**. Trials were selected based on the strength of correlation with each participant’s accuracy-maximizing principal component, corresponding to MI test accuracy. EEG activity was measured during the temporal error monitoring period (between response onset and offset). Channel selection frequency peaked at POz, P6, PO4, P4, P8, PO8, AFz, AF8, CP6, and CP2, revealing distributed parieto-frontal contributions to single-trial MI decoding. **b Cross-frequency integration maximizes MI decoding accuracy**. Each dot represents participant-wise, fold-averaged test accuracy. Decoding exceeded chance level (50%) for all frequency bands combined, as well as for θ, α, and β individually. Combined-band accuracy was higher than that of α or β alone, but not significantly higher than θ, suggesting a prominent contribution of θ-band activity to metacognitive processing and indicating that integrating information across frequency bands maximizes the extraction of higher-order evaluative signals. **c–f Temporal dynamics of metacognitive signals across task phases**. Classification accuracy is compared between the TP onset (light gray) and the TEM decision phase (dark gray). Accuracy was higher during the TEM decision phase across all frequency bands, indicating that metacognitive signals strengthen during explicit error evaluation. However, combined-band activity showed above-chance MI decoding already at TP onset, suggesting that evaluative neural markers are present before explicit decision-making. α-band activity exhibited a trend-level effect at TP onset, indicating that specific oscillatory components may contribute to early metacognitive evaluation. Overall, these findings show that metacognitive signals peak during the decision phase but remain partially detectable during ongoing action timing, suggesting that TEM and higher-order evaluation unfold continuously rather than being confined to a discrete decision point.

Comparisons between TP onset and the TEM decision phase revealed clear temporal specificity in the emergence of MI-related signals (Figure 5c–f). At TP onset, *θ*- and *β*-band decoding did not exceed chance, but both became significant during the TEM decision phase (*θ*: *t*(28) = −3.911, *p* < .001; *β*: *t*(28) = −3.221, *p* = .002). *α*-band decoding showed only a trend-level effect at TP onset but reached significance during the decision phase (*t*(28) = −2.429, *p* = .019). Notably, combined-band decoding (θ+α+β) was already above chance at TP onset and increased further during the TEM decision phase (*t*(28) = −2.707, *p* = .009), indicating that cross-frequency integration provided the strongest and most sustained metacognitive signal across both task phases.

Overall, these results indicate that although MI-related neural signals were detectable during action execution, their predictive strength peaked during the TEM decision phase. This pattern of temporally graded emergence is consistent with a continuously accumulating higher-order evaluative process, in which metacognitive information becomes maximally accessible immediately preceding explicit error judgment.

## 7) Discussion

The present study investigated whether metacognitive evaluation of motor action timing can be explained by the same neural signals that support timing performance, or whether it reflects at least partially distinct processes involving additional evaluative computations. To adjudicate between these accounts, we applied Vision Transformer decoding to single-trial EEG recorded across θ, α, and β frequency bands during both the motor action execution and temporal error monitoring (TEM) phases. The findings converge on a consistent picture: motor action timing and metacognitive evaluation are supported by partially dissociable neural architectures, with the latter requiring a form of multi-band integration that the former does not.

Time production categories were reliably decodable from any individual frequency band. θ, α, and β each contributed sufficient information for classification, and broadband combination yielded no substantial improvement. This pattern of partially redundant, band-sufficient encoding is consistent with prior evidence that β-band oscillations index temporal intervals at the single-trial level,^17,18^ and propose that frequency-specific oscillatory codes provide parallel, largely independent representations of interval duration. The selective contribution of β and θ to TP decoding, and the relative weakness of α, aligns with the established topography of post-movement β rebound^4,17,18^ and with fronto-parietal θ coordination during motor action and decision processes.^37,38^

In contrast, metacognitive inference (MI), the trial-by-trial alignment between objective timing performance and subjective error evaluation, was decodable only when θ, α, and β were integrated simultaneously. No individual band supported reliable MI classification. This dissociation is not merely a quantitative difference in decoding strength; it reflects a qualitative difference in representational architecture. Single-process accounts^12,16^ predict that metacognitive evaluation should be decodable from the same signals as timing performance, since both arise from a common accumulator variable. The data directly contradict this prediction. The failure of individual bands to support MI decoding, combined with the success of their joint integration, implies that metacognitive evaluation draws on coordinated multi-band dynamics that are not reducible to any single oscillatory substrate. This is consistent with the Higher-Order Representation (HOR) framework,^14,15^ which holds that conscious awareness of a mental state requires a higher-order re-representation that takes the first-order state as its object rather than simply reflecting it.

The role of β within this multi-band context deserves particular attention. β participated in MI decoding only as part of the integrated broadband signal, not as an independent predictor, a pattern distinct from its role in TP decoding, where it was sufficient alone. This dissociation is consistent with proposals that β reflects the maintenance and reactivation of internal representational states beyond simple motor execution feedback,^39,40^ and with direct evidence that post-movement β indexes confidence in internal model estimates in sensorimotor contexts.^41^ The present findings extend this interpretation to the domain of temporal metacognition: β contributes to metacognitive evaluation not as a motor readout but as part of a coordinated higher-order representation that also requires θ- and α-band contributions.

Importantly, MI was already decodable during ongoing motor execution, up to 1 s before action termination, and this decoding was strengthened further during the explicit TEM decision phase. This temporal profile indicates that evaluative information accumulates continuously during interval production rather than arising solely at the point of explicit judgment. This finding aligns with Bayesian accounts of temporal uncertainty,^42,43^ under which uncertainty about the current production is represented as a structured signal that accumulates progressively during interval generation and is available for metacognitive evaluation prior to any explicit response. It is also consistent with continuous evidence accumulation frameworks in which evaluative signals build progressively in parallel with ongoing performance, rather than emerging only after task completion.^44,45^

Spatial evidence further supported this dissociation. TP decoding localized to centro - parietal and frontal-central regions, consistent with the established topography of motor action timing and post-movement β rebound. MI decoding, by contrast, recruited a distributed parieto-frontal network spanning posterior and prefrontal sites, consistent with the integration of temporal performance representations and prefrontal evaluative processes.^46^ This spatial shift from execution-related to evaluative networks is consistent with neural evidence that dedicated prefrontal circuits contribute to higher-order representations in ways that cannot be reduced to first-order activity.^3,15^

The most direct evidence against single-process accounts came from the neural– behavioral dissociation. Participants stratified by TP decoding accuracy showed no difference in error monitoring strength or metacognitive distance, directly contradicting the prediction that stronger timing encoding should produce more accurate self-evaluation. Participants stratified by MI decoding accuracy, by contrast, showed significantly stronger performance–error response correlations and reduced metacognitive distance, despite equivalent timing accuracy. Because timing performance was equated across groups, the metacognitive benefit cannot be attributed to superior timing decoding; it must arise from a representational layer that is neurally distinct from performance control. This selective pattern challenges hierarchical accounts as much as single-process ones: even a model that posits a distinct metacognitive stage but ties its precision to the quality of first-order (here, motor timing signals) encoding^13^ would predict some relationship between TP decoding accuracy and TEM precision, an absent relationship. The data are instead most consistent with HOR theory,^14,15^ under which metacognitive evaluation constitutes an active re-representation that draws on first-order timing states but engages at least partially distinct mechanisms beyond those supporting task execution.

TEM confidence judgments showed a pattern distinct from both TP and MI decoding. Significantly above-chance decoding was observed for α and β individually, without requiring multi-band integration. This suggests that subjective certainty may partly reflect first-order signal properties, β-indexed performance quality and α-indexed attentional certainty, rather than the higher-order inferential process that underlies accurate metacognitive evaluation. The dissociation between confidence and MI decoding raises the possibility that these two aspects of metacognition, the accuracy of self-evaluation and the subjective sense of certainty accompanying it, are at least partially separable at the neural level, a distinction that aligns with signal detection frameworks separating metacognitive sensitivity from response bias.^47,48^ Taken together, these findings establish multi-band neural integration as a necessary condition for temporal metacognition. The dissociation between single-band sufficiency for timing performance and multi-band dependence for metacognitive inference, combined with the selective behavioral relevance of MI decoding accuracy, converges on the conclusion that metacognitive evaluation is not a passive reflection of interval production signals but an active, higher-order process that recruits a representational organization qualitatively distinct from the one supporting primary task performance. Within the broader landscape of temporal metacognition,^4^ this constitutes the first single-trial neural evidence directly distinguishing first-order timing representations from second-order metacognitive ones, and provides direct support for the view that self-evaluative processing requires a dedicated neural architecture consistent with HOR theory.

## 8) Limitations and Future Directions

Several methodological considerations merit acknowledgement. First, trial–channel observations per participant after preprocessing were modest. Replication with larger trial samples and multi-session designs may represent an important future step. Future work might also benefit from data augmentation techniques to expand single-trial training sets,^49^ and from study designs incorporating subject-specific EEG architectures to better accommodate inter-participant variability.^50^

Second, isolating metacognitive signals at the single-trial level is inherently challenging: the brain may not always produce clearly detectable β-rebound-like patterns, and metacognitive signals may be particularly difficult to decode reliably. To address this, we developed a PCA-based pipeline retaining the top 30% of oscillatory-structure-aligned trials, a step that may have been critical for reliable decoding and could offer a useful methodological contribution for future single-trial metacognitive EEG studies.

Third, recording under a single motor condition leaves open whether the θ–α–β integration pattern identified here might be disrupted or reorganized under higher motor demands. Given that increased motor demands selectively impair TEM precision without affecting timing accuracy,^22^ future work combining parametric motor manipulations with single-trial EEG decoding could potentially provide the mechanistic bridge needed to connect the behavioral and neural findings of this thesis.

## Acknowledgements

This research was supported by the National Science Centre, Poland (grant no. 2019/35/B/HS6/04389 to TWK). We thank Hatice Balkan for assistance with EEG data collection. We also thank Izabela Szumska and Dorota Stelmaszyńska for technical assistance in the experimental setup.

## 9) Code and Data Availability

The data and code supporting the findings of this study will be made openly available on GitHub at https://github.com/TEM-Opus2020 upon publication.

