## Supplemental Table 1 for "The Brain as Actor and Evaluator: Distinct Neural Codes for Timing and Self-Evaluation of Timing Errors Revealed by Transformer-Based Decoding"

| | | | | |
|---|---|---|---|---|
| **Session 1** | EEG + tDCS Set up | **Block:** tDCS - EEG | Break | **Block**: EEG-only |
| **Session 2** | EEG + tDCS Set up | **Block:** Sham - EEG | Break | **Block**: EEG-only |

**Table 1. EEG Experiment Session Structure.** Table 1 summarizes the structure and order of experimental sessions completed by participants. Each participant completed two sessions on separate days, with the order of sessions counterbalanced across participants (indicated by the blue arrow). The present study analyzed EEG-only data obtained from the Sham condition.
